# Simultaneous transcriptome and methylome profiles of single mouse oocytes provide novel insights on maturation and aging

**DOI:** 10.1101/2020.08.22.260612

**Authors:** Yan Qian, Qin Cao, Jinyue Liao, Chun Shui Luk, Ashley Hoi Ching Suen, Annie Wing Tung Lee, Ting Hei Thomas Chan, Judy Kin Wing Ng, Nelson Leung Sang Tang, Hoi Sze Chung, King Lau Chow, Tak Yeung Leung, Ching-Hung Chen, Wen-Jui Yang, Jack Yu Jen Huang, Wai-Yee Chan, David Yiu Leung Chan, Tin Chiu Li, Kevin Y. Yip, Tin-Lap Lee

**Affiliations:** Developmental and Regenerative Biology Program, School of Biomedical Sciences, The Chinese University of Hong Kong, Shatin, N.T., Hong Kong SAR, China; Department of Computer Science and Engineering, The Chinese University of Hong Kong, Shatin, N.T., Hong Kong SAR, China; Department of Chemical Pathology, The Chinese University of Hong Kong, Shatin, N.T., Hong Kong SAR, China; Department of Obstetrics and Gynaecology, The Chinese University of Hong Kong, Prince of Wales Hospital, Shatin, N.T., Hong Kong SAR, China; Division of Life Science, Hong Kong University of Science and Technology, Hong Kong SAR, China; Department of Fertility and Reproductive Medicine, Ton-Yen General Hospital, Hsinchu County, Taiwan; Institute of Molecular and Cellular Biology, National Tsing Hua University, Hsinchu, Taiwan; Division of Infertility and Reproductive Medicine, Taiwan IVF Group Centre, Hsinchu City, Taiwan

**Keywords:** Oocyte, Advanced maternal aging, Single cell, Transcription, DNA methylation, Quality prediction model

## Abstract

**Background:** Advanced maternal aging has become a worldwide public health issue that contributes to female fertility decline and significant risk to embryo development. Despite transcriptional and epigenetic alterations reported in oocyte maturation and development, the dynamics of gene expression and DNA dynamics associated with aging remain elusive. Here we generated simultaneous transcriptome and methylome profiles of mouse oocytes during aging and maturation at single-cell and single-base resolution to examine key biological processes and identify the key targets for novel treatment options.

**Results:** We report the dynamics in transcriptome and DNA methylome in mouse oocytes during maternal aging and oocyte maturation. Age-associated gene expression changes showed mitochondrial dysfunction in GV oocytes and defects of chromosome segregation and spindle assembly in MII oocytes. EIF2 signaling protein synthesis pathway was also impaired during aged oocyte maturation. Moreover, distinctive DNA methylation patterns were demonstrated during maternal aging in GV and MII oocytes. A positive correlation between gene expression and methylation in gene body was characterized. Furthermore, we identified several promising biomarkers, including IL-7, to assess oocyte quality, which are potential therapeutic targets for improve oocyte maturation. More importantly, we built the first mouse oocyte maturation and age prediction model using transcriptome data and validated its feasibility in published data.

**Conclusions:** This work provides a better understanding of molecular and cellular mechanisms during mouse oocyte aging, points a new direction of oocyte quality assessment, and paves the way for developing novel treatments to improve oocyte maturation and quality in the future.

## Background

Maternal age-related fertility decline is resulted from several mechanisms, including the dynamics in reproductive hormones, the apoptosis of granulosa cells in the ovarian follicle and age-dependent inflammation in reproductive organs [1]. However, oocyte *per se*, as the source for fertility, is not understood precisely in the process of maternal aging, possibly due to the limitation of rare experimental materials and sophisticated experimental technologies.

Cellular dynamics has been investigated extensively in oocytes with maternal aging. Aged oocytes demonstrate various mitochondrial defects [2]. Mitochondrial swelling, reduced ATP and metabolic activity, decreased number of mitochondria and impairment in mitochondrial-DNA (mtDNA) repair systems have been reported as contributors to poor quality of aged oocytes [3–5]. Besides, telomeres, which are essential in meiosis, are reported to be shorter in aged oocytes. This could in turn promote genomic instability, apoptosis and cell cycle arrest [6]. However, molecular alterations, including those at the DNA, RNA, and epigenetic levels, during oocyte aging have not been comprehensively investigated.

In recent years, there have been growing interests in molecular changes during human aging and senescence. Multiple studies have demonstrated that transcriptome profiles in various tissues, including lung, kidney, brain and peripheral blood cells, are altered during aging [7–10]. The difference between biological age and chronological age has been revealed in human dermal fibroblasts [11]. Besides transcriptome, genome variant acquirements have been reported during human aging [12]. Also, accumulation of aging-related copy number variations (CNVs) has been displayed in aged human brain [13]. Furthermore, aging-associated epigenetic modifications, especially DNA methylation, have also received much attention in the past decades [14]. Horvath established the first DNA methylation clock to predict biological age of multiple human tissues based on the methylation dynamics of 353 CpG sites [15]. In the same year, Hannum G et al. built a DNA methylation clock for single tissues [16]. However, to date, no similar molecular markers or prediction tools have emerged for oocytes.

Single-cell sequencing technologies have been developed rapidly. Single-cell profiles of the genome, transcriptome, methylome, and histone modifications have been generated by various single-cell sequencing technologies. These single-cell approaches provide a better understanding of cell heterogeneity, which allow us to identify novel cell populations. However, the regulatory mechanisms of gene expression could not be illustrated directly and precisely in individual cells based on transcriptome profiles alone. To solve this issue, Angermueller et al. firstly integrated the methods of single-cell genome sequencing and single-cell transcriptome sequencing to study the diverse effects of genetic variations on gene expression in the same single cells [17]. Later, Angermueller et al. reported another method for parallel sequencing of methylome and transcriptome on individual cells [18]. These parallel sequencing technologies offer us chances to obtain as much molecular information as possible from a single cell and to gain a complete understanding of the regulatory mechanisms among the genome, transcriptome, and methylome.

In this study, we aim to investigate the dynamics of the transcriptome and methylome of mouse oocytes during maternal aging and characterize aging-related molecular signatures in mouse oocytes to predict oocyte quality. We applied a modified version of scM&T-seq to generate simultaneous single-cell transcriptome and methylome profiles of single mouse mature (MII) and immature (GV) oocytes at different ages (6 weeks old & 12 months old). We identified transcripts with significantly altered expression and differential methylated regions (DMRs) and demonstrated various molecular mechanisms, especially the methylation dynamics of non-CpG sites, contributing to oocyte aging. Besides, we used DNA methylation data to model gene. expression. Based on the transcriptome dynamics during oocyte aging and maturation, we identified aging-related and maturation-related gene signatures and established two prediction models for mouse oocyte age and maturation respectively. To the best of our knowledge, our study is the first to employ single-cell sequencing technologies to explore molecular alterations and regulatory mechanisms in mouse oocytes during maternal aging. Our prediction models predicted the molecular age and maturation stage of mouse oocytes with good accuracy and the same method can be applied to develop analogous models for human oocytes as a complement to the current morphology assessment approach in the clinic.

## Results

### Parallel transcriptome and methylome capture on immature and mature mouse oocyte aging models

To investigate the effects of age on gene expression and DNA methylation, we harvested GV oocytes and MII oocytes from 6-week old (young) and 12-month old (aged) female C57BL/6 mice. The 12-month time point was selected since the fertility of female mice dramatically declines after 12 months (Book <u>Mouse genetics: concepts and applications</u>.). The four groups are referred to as GV-6wk, GV-12m, MII-6wk and MII-12m hereinafter, with six oocytes in each group (Figure 1).

**Figure 1.**
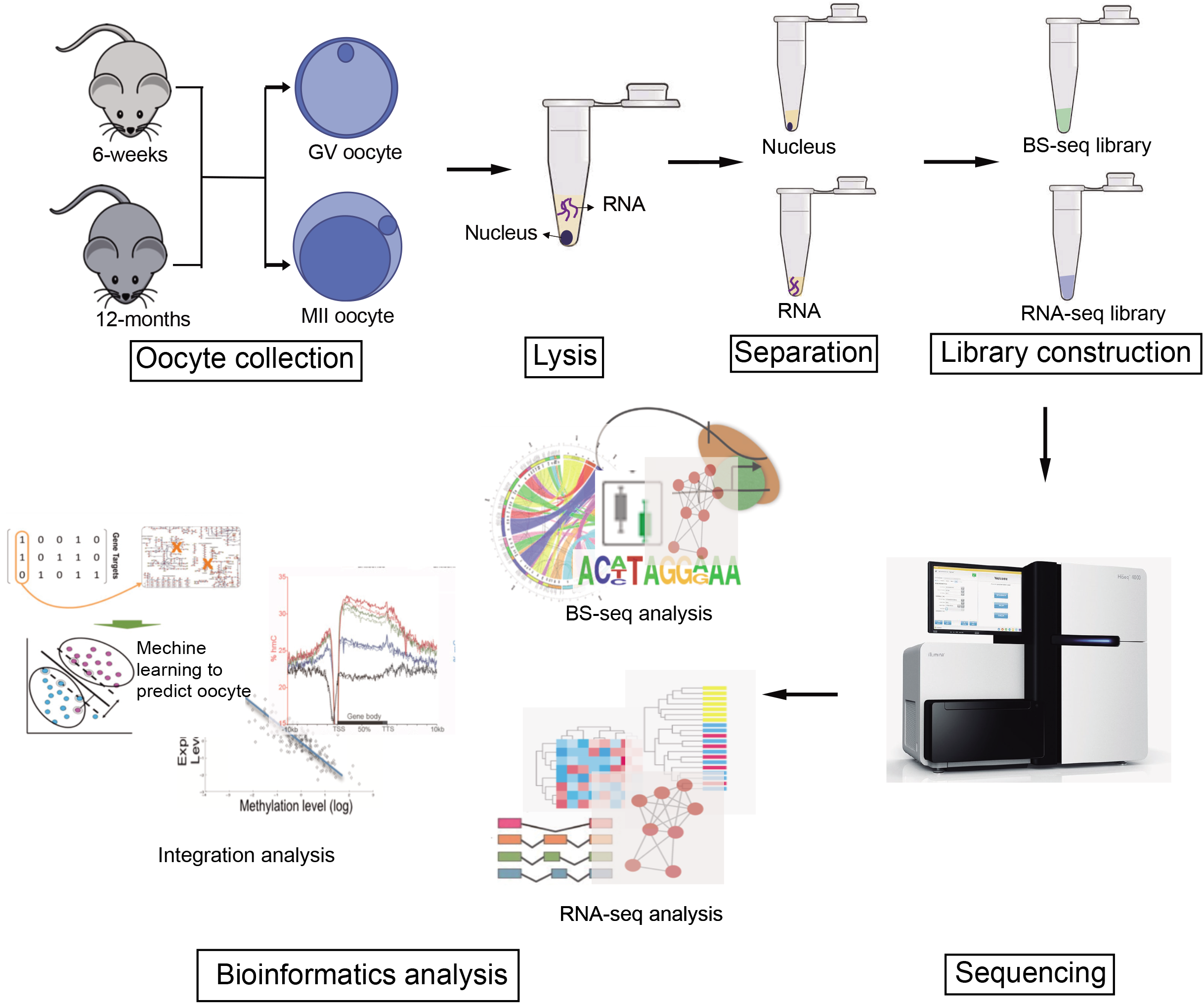
Schematic diagram of study. In this study, we collected both GV and MII oocytes from C57/BL6 female mice at the age of 6 weeks and 12 months, and assigned them to the groups of GV-6wk, GV-12m, MII-6wk and MII-12m. For each oocyte, Soft buffer is used to release RNA in cytoplasm without destroying the nucleus. The cytoplasmic RNA was subjected to transcriptome sequencing by SMART-seq2 method, and the intact nucleus was used to perform single-cell bisulfite sequencing (scBS-seq). Analysis of the transcriptome, DNA methylome, and their integration was performed, and a model to predict oocyte quality was established.

We modified scM&T-seq by performing whole genome bisulfite sequencing (WBS) instead of previous reduced representation bisulfite sequencing (RRBS) on separated DNA to obtain more comprehensive DNA methylation information. We applied the modified scM&T-seq on individual oocytes to separate DNA and RNA from the same oocyte. For transcriptome profiles, SMART-seq2 was performed on the released RNA from each of the six individual oocytes in each group, resulting in 7.21M ± 2.73M raw reads obtained from each library. The average mapping efficiency was 57.38±11.67%, and the multiple alignment rate was 7.35±4.54%. After alignment to the reference genome (mm10) with the annotation set of Ensembl GRCm38.p5, we detected 18203 ± 3377 transcripts in each sample (Additional file 2: Table S1). Principal component analysis (PCA) showed that different groups clustered separately, though samples in the MII-12m group were most dispersed (Figure 2a), which indicates that MII oocytes had higher heterogeneity in transcriptome than GV oocytes. Pairwise Pearson’s correlation analysis demonstrated that oocytes in the same group had much higher correlation coefficients, and the aggregated gene expression profile from individual cells of a group recapitulated the individual cells’ profiles well (Additional file 1: Figure S1a & Figure 2b). Furthermore, oocyte-specific genes, such as *Gdf9, Dppa3*, and *Zp1*, were highly expressed in all the samples, while the markers for cumulus cells, such as *Cd14* and *Fshr*, were hardly detectable (Figure 2c). This suggested that our oocyte samples were not contaminated by cumulus cells. Furthermore, several genes were picked to validate their expression level in oocytes by real-time PCR, and the results showed the consistent trend of gene expression during aging and maturation with the RNA-seq results, which confirmed the reliability of the sequencing data (Additional file 1: Figure S1).

**Figure 2.**
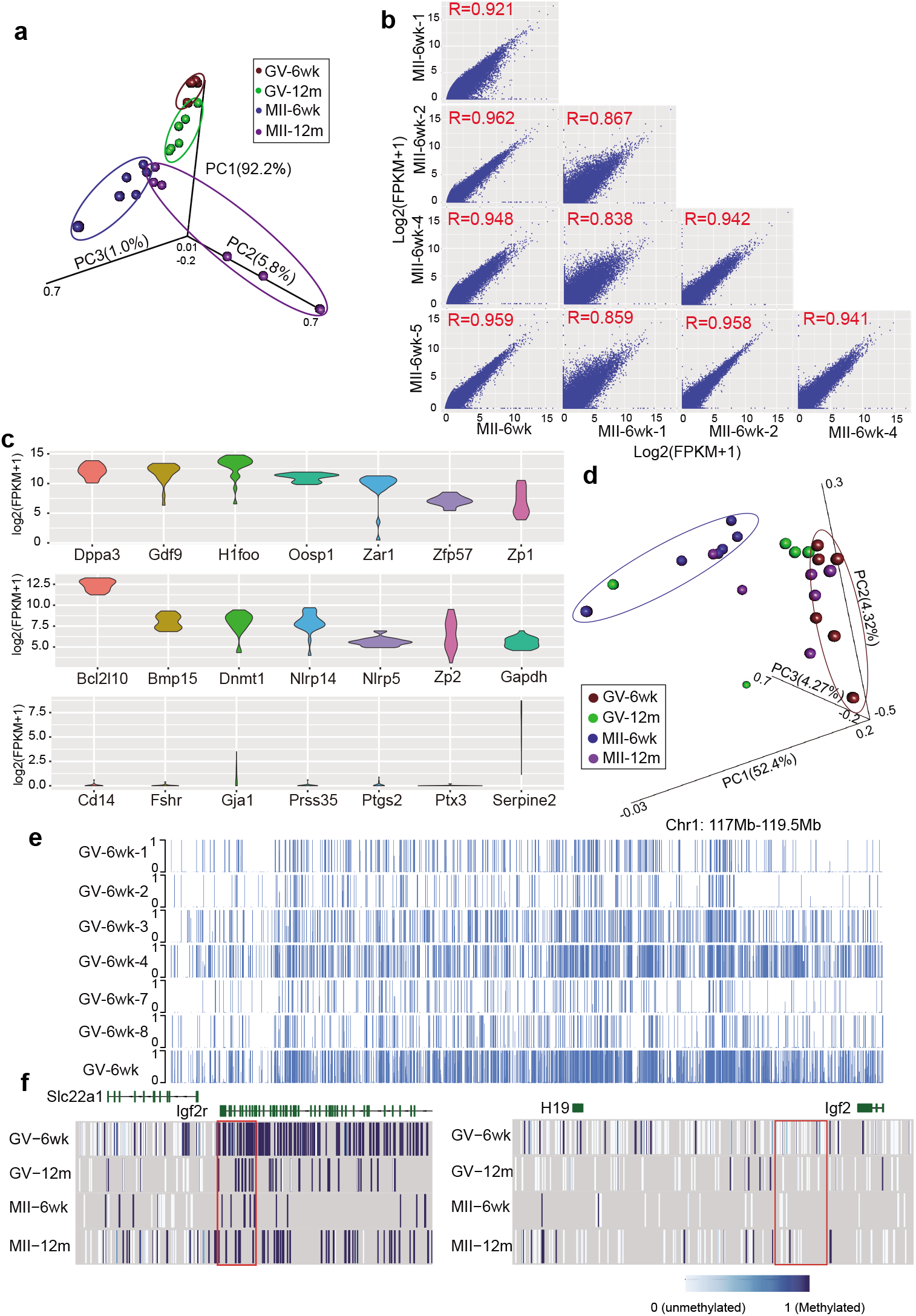
scM&T-seq on mouse oocytes. **(a)** Principal component analysis (PCA) on 24 oocytes in 4 groups (GV-6wk: brown; GV-12m: green; MII-6wk: blue; MII-12m: purple) based on genes detected in all samples. **(b)** Pairwise correlation comparison of gene expression levels (log2FPKM+1) for 4 randomly picked individual MII-6wk oocytes and the MII-6wk group average. **(c)** Violin plots showing the substantial expression variability of oocyte specific-genes (Top and Middle panel) and cumulus cell associated genes (Bottom panel). **(d)** PCA on 23 oocytes in 4 groups, excluding one GV-12m oocyte with the lowest mapping efficiency, based on the mean CpG methylation level of gene bodies detected in all sample. CpG sites covered with less than three reads and gene bodies with less than 5 CpG sites are filtered. The gene body is defined as the region from the transcription start site (TSS) to the transcription terminal site (TTS). **(e)** Line plots demonstrating CpG methylation level in a random region of 2057kb from scBS-seq data of individual GV-6wk oocytes (Top 6 tracks) and in silico pooled GV-6wk group data (Bottom track). **(f)** Heatmap showing the methylation profile of CpG sites in *Igf2r* (Left), a maternally imprinted gene, and *H19/Igf2* region, a paternally imprinted control region, for four in silico-merged groups scBS-seq datasets. The imprinted regions are highlighted in red frames.

To generate single-cell methylome profiles of oocytes, we performed single-cell bisulfite sequencing (scBS-seq) on each nucleus of oocyte and obtained an average of 15.42M ± 6.25M paired reads per cell (Additional file 2: Table S2). The bisulfite conversion efficiency ranged from 97.121% to 99.062%, comparable to other scM&T-seq or scBS-seq studies [18, 19]. After alignment to the reference genome (mm10), the mean total mapping efficiency was 36.93%, while one sample got an extremely low mapping efficiency with less than 5% (4.21%) and was excluded in the following analysis. We detected an average of 1.17M CpG sites, 5.11M CHG sites, and 17.40M CHH sites in each sample. We observed an unexpected high duplication rates of sequencing reads, from 56.64% to 99.11%. This indicated the potential of over-amplification during library construction, the protocol of which deserves better optimization in the future. It would be better to look for better bisulfite conversion protocol to reduce DNA lose or employ less cycle numbers of PCR amplification to reduce high duplication rates. To examine the quality of our scBS-seq data, we compared the data resolution of individual oocytes with the merged data of oocytes from the same group, and it showed high repeatability of single cell data (Figure 2e). Also, CpG sites in the known maternally imprinted region, *Igf2r*, were fully methylated. Meanwhile, those in the paternally imprinted region, *H19/Igf2*, were nearly unmethylated (Figure 2f). All of these results confirmed the success in generating simultaneous transcriptome and methylome profiles of single oocytes by scM&T-seq.

### Decoding transcriptional dynamics during oocyte aging

To explore the effect of aging on the transcriptome of oocytes at GV stage, we identified 1377 differentially expression genes (DEGs) during GV oocyte aging. Of them, 631 genes was downregulated during aging, while the remaining 745 genes were upregulated. Gene ontology (GO) analysis showed that the downregulated genes were enriched in RNA processing, mitochondrion organization, and ATP synthesis, while the enriched functions of upregulated genes were cell cycle, DNA repair and chromosome organization (Figure 3a). The decline of mtDNA copy numbers, mtDNA deletion, and reduced production of coenzyme Q were reported in aging oocytes, while defects in mitochondrial function affect oocyte maturation [2]. This suggested that the transcription dynamics of genes in mitochondria related function during GV oocyte aging would contribute to mitochondrial dysfunction, which plays a critical role in the reduced oocyte quality.

**Figure 3.**
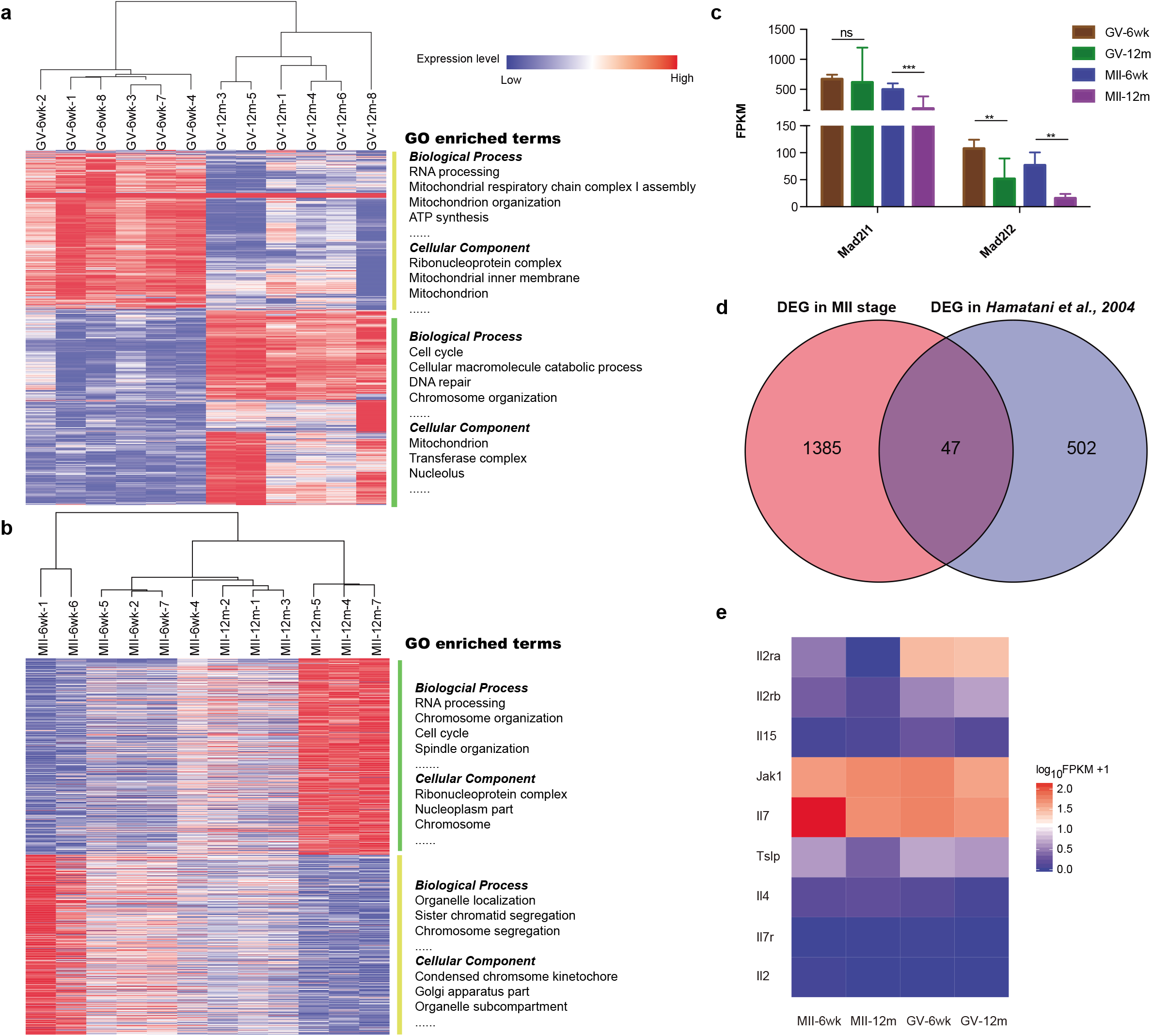
Age affecting expression of different functional genes in GV and MII stage. **(a-b)** Heatmap showing differentially expressed genes (DEGs) during GV oocyte aging (a) and MII oocyte aging (b). Oocytes in GV-6wk and GV-12m are clustered based on the expression difference of these 1377 genes (a) and 1432 genes (b). GO enrichment analysis results on upregulated DEGs (green), and downregulated DEGs (yellow) are listed in terms of Biological Process and Cellular Components. **(c)** Bar plots demonstrating the expression level of two *Mad2* genes (*Mad2l1* and *Mad2l2*) in four groups (GV-6wk: brown; GV-12m: green; MII-6wk: blue; MII-12m: purple). The comparisons are performed with One-way ANOVA (ns: not significant; *p<0.05; **p<0.01; ***p<0.001). **(d)** Comparison of age-associated DEGs in MII stage between our data and published data [29]. A total of 47 genes were common in two gene sets. **(e)** Heatmap illustrating expression level (log10FPKM+1) of genes involved in Il-7 pathway in 4 groups.

Furthermore, we employed Ingenuity Pathway Analysis (IPA) to perform the upstream analysis on the DEGs to predict the potential activation state of upstream regulators. The active states of lipopolysaccharide, tumor necrosis factor (TNF), and phorbol myristate acetate (PMA) were predicted in the aged oocytes based on the expression changes of their downstream genes in our dataset (Additional file 1: Figure S2). Exposure to lipopolysaccharide was reported to inhibit the maturation potential of bovine oocytes by affecting cell cycle, cytoskeletal dynamics, oxidative stress, and epigenetic modifications [20]. In porcine and cattle, TNF could inhibit the maturation of oocytes from GV to MII stage and increase the proportion of oocytes with abnormal chromosome alignment and cytoskeleton structure [21, 22]. On the other hand, PMA could inhibit the meiotic maturation in mouse oocyte [23]. Combined with previous results, our data suggested that maternal aging would have similar effects on the transcriptome of GV oocytes as these upstream regulators, hence reducing the maturation potential of oocytes through mechanisms, including cell cycle, oxidative stress and mitochondrial dysfunction.

Next, we compared the transcriptome of MII oocytes at different ages and identified 1432 DEGs, including 706 upregulated DEGs and 726 downregulated DEGs. GO enrichment analysis showed that these upregulated genes were enriched in RNA processing, chromosome organization, cell cycle, and spindle organization, while GO terms including organelle localization, sister chromatid segregation and chromosome segregation were obtained from these downregulated genes (Figure 3b). Previous studies reported that the meiotic spindle was frequently found to be abnormal in aged human oocytes [24], as spindles became elongated and smaller, and fewer microtubular foci were detectable [25]. Our results showed that the expression of the spindle checkpoint protein gene, *Mad2*, was significantly decreased in aged MII oocytes (Figure 3c), which was consistent with previous studies [26, 27]. Different from GV oocytes, mitochondria-related function was not enriched in MII oocytes. This suggested that mitochondrial dysfunction often occurred in GV oocytes while the defects in chromosome segregation and organization were more pronounced in MII oocytes during maternal aging. Moreover, upstream regulator analysis by IPA found that *Kdm5b*, an histone H3 lysine 4 (H3K4) demethylase, was functionally inhibited based on aging associated DEGs in MII oocytes (Additional file 1: Figure S3). This was consistent with the previous finding that genome-wide H3K4 dimethylation (H3K4me2) was increased in older mouse MII oocytes [28].

Moreover, we compared our data with published microarray data of aging mouse MII oocytes [29]. A total of 47 DEGs were shared in two datasets (Figure 3d). Interleukin 7 (IL-7), a maturation-specific oocyte-secreted factor, was among them. At both MII and GV stages, *Il-7* was downregulated during aging (Figure 3e). Further investigating on genes involved in the IL-7 pathway, we found *Jak1* (Janus Kinase 1) was expressed highly but stably in all four groups, but *Il-2ra* (Interleukin receptor 2a) showed a high expression level in GV stage and differential expressions during aging of both GV and MII oocytes (Figure 3e), suggesting that *Il-2ra* might also be involved in oocyte aging.

### Demonstrating dynamics of gene expression during oocyte maturation

Besides exploring the age effect on the transcriptome of oocytes at individual stages, we asked whether age affected the changes in transcriptome that normally occur during oocyte maturation. We first identified 4613 maturation-related DEGs in six weeks old and 4223 maturation-related DEGs in 12 months old mice (Figure 4a). The relative changes of maturation (fold change) were compared between ages of 6 weeks and 12 months. These DEGs showed high Pearson’s correlation (r = 0.8181), suggesting that most of them shared the same maturational expression pattern. To characterize DEGs in which maturational expression changes were affected by age, we established a linear regression model and used a statistical cutoff (95% prediction interval) to define 260 DEGs (Figure 4b). These 260 genes had different levels of maturation-related differential expression at 6 weeks and 12 months, and they were found responsible for functions in the metabolic processes (Figure 4d). Several reproductive processes or sexual reproduction-related genes, such as *Mael* (Maelstrom spermatogenic transposon silencer), *Sebox* (Skin-, embryo-, brain- and oocyte-specific homeobox) and *Elf2* (Eukaryotic translation initiation factor 2), were included in these DEGs (Figure 4c). Meanwhile, canonical pathway analysis on these DEGs demonstrated that the EIF2 pathway, a critical protein synthesis pathway, was the most significantly enriched canonical pathway (Figure 4d). EIF2 regulation pathway and regulation of eIF4 and p70S6K, which were also enriched in the canonical pathway analysis, were reported to play an essential role in degrading transcripts in GV-MII transition [30]. Our data suggested that maternal age might disrupt the regulation of protein synthesis during oocyte maturation through EIF2/EIF4 related pathways.

**Figure 4.**
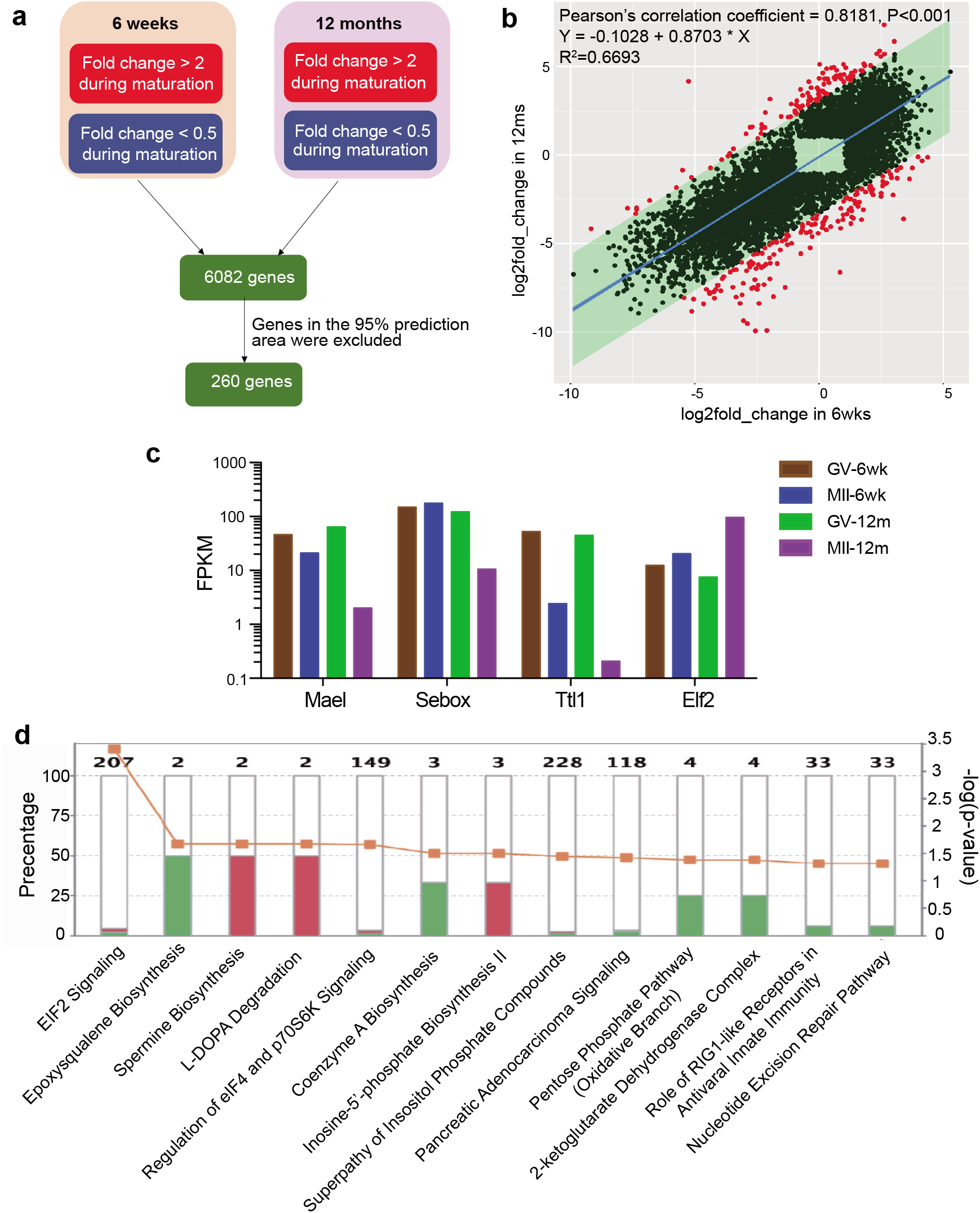
Age destroying the normal changes of transcriptome during oocyte maturation. **(a)** The strategy of identifying genes whose expression changes during maturation are affected by maternal age. **(b)** Scatter plot demonstrating maturation associated expression changes of 6082 genes at two ages. Fold-change (in log2 scale) of each gene during maturation in 12m were plotted against that in 6wk. The regression model is calculated and displayed as the blue line. The 95% prediction interval, which is calculated by the expression changes of genes in 6 weeks as variables, is displayed as a green region. The red dots show genes whose actual maturation associated changes at 12 months old exceed the 95% prediction interval. **(c)** Bar charts showing the mean expression pattern of reproduction-related genes involved in 260 identified genes. **(d)** Canonical pathway analysis on 260 identified genes by Ingenuity Pathway Analysis (IPA). Red and green bars represent genes with up- and down-regulation during maturation respectively. The number above each bar indicates the number of genes involved in the corresponding pathway and the orange dots display the enrichment scores scaled by the right y-axis.

Isoform switch has been implicated in multiple physiological and pathological processes, such as embryonic development and cancers (Ref PMID:26939613, 27916660). It has also been observed in aging and aging-related diseases [31]. Thus, we wondered whether isoform switch would occur during oocyte aging. We identified 56 age-related isoform switches in GV oocytes and 377 in MII oocytes by IsoformSwitchR [32]. The dominant biological mechanism of age-related isoform switches was alternative splicing (Figure 5a-b). Genes with age-related isoform switch in GV oocytes showed functional enrichment in organelle fission, nuclear division, and chromosome segregation. Meanwhile, regulation of cell cycle, DNA repair and mRNA processing were enriched in age-related isoform switch in MII oocytes (Figure 5c-d). Importantly, we found that a critical *de novo* methyltransferase, *Dnmt3a*, demonstrated age-dependent isoform switch in GV oocytes. The coding isoform 2, *Dnmt3a2*, made the most contribution to the increased expression of *Dnmt3a* during GV oocyte aging (Figure 5e), which suggested that maternal age influenced *de novo* methylation in GV oocytes mainly by *Dnmt3a2* instead of other isoforms. On the other hand, the abundance of a longer and coding isoform of *Txnip* (thioredoxin-interacting protein), an essential regulator of glucose metabolism and cytoplasmic stream, was observed in aged GV oocytes (Figure 5f), which suggested a higher protein level of *TXNIP* in aged GV oocytes even though its total RNA level was relatively stable during GV oocyte aging.

**Figure 5.**
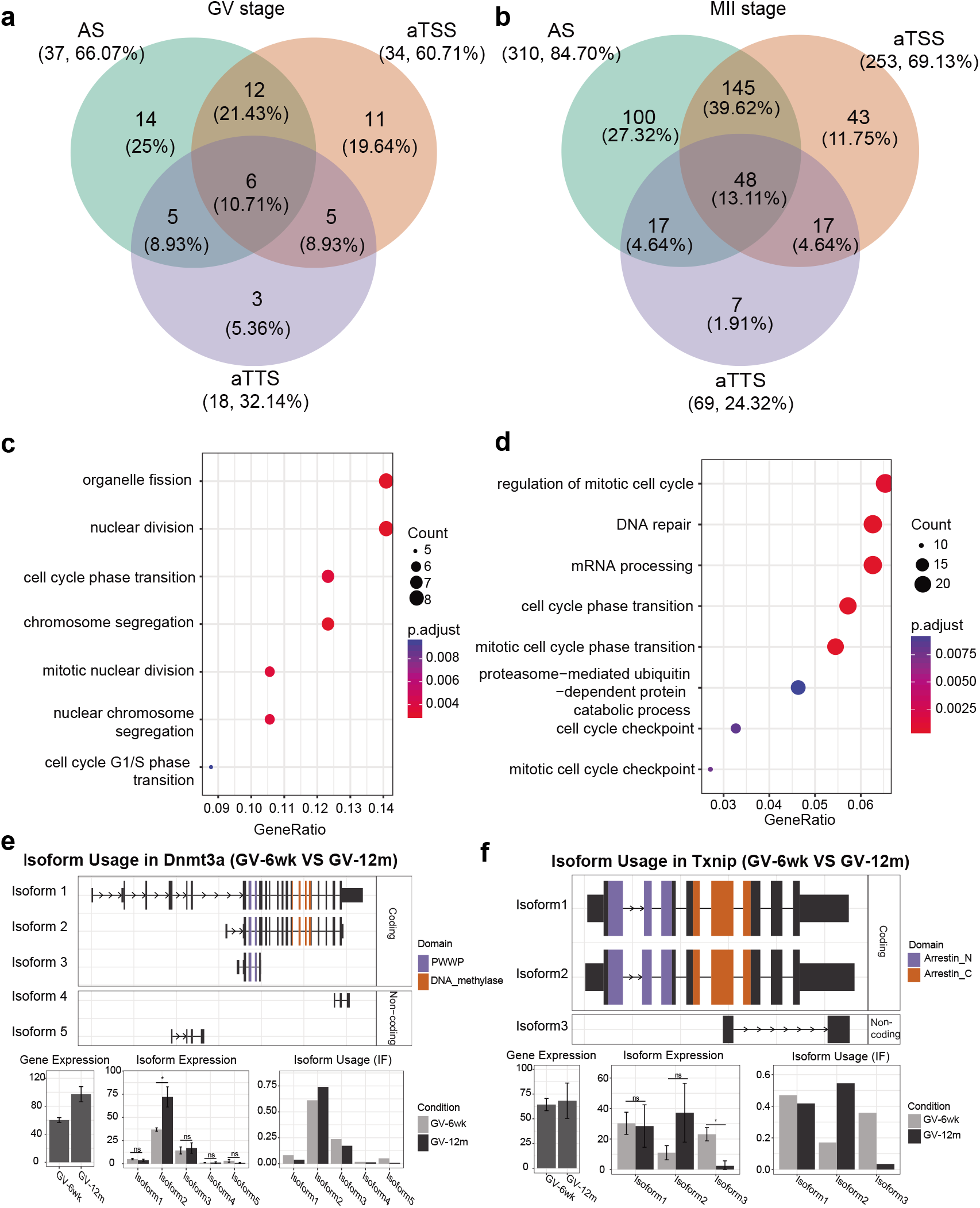
Maternal age contributing to isoform switches. **(a-b)** Venn diagrams showing the numbers of different biological mechanisms for age-related isoform switches in GV (a) and MII (b) oocytes (AS: alternative splicing; aTSS: alternative transcription start site; aTTS: alternative transcription terminal site). **(c-d)** GO analysis on genes with identified age-related isoform switches in GV (c) and MII (d) oocytes. The dot size represents gene count, and the color of dots shows adjusted p values. **(e-f)** Gene examples of identified age-related isoform switches, Dnmt3a (e) and Tnxip (f). The upper panel displays the structures of different isoforms, corresponding functions and important domains. The lower panel shows the gene expression of the gene in two groups (Left), the isoform expression (Middle) and the contribution of each isoform to the total expression of the gene (Right). (ns: not significant; *p<0.05)

### Revealing age-associated changes in methylome

We then proceeded to investigate the effect of age on methylome in oocytes. During the analysis of methylome, we excluded one sample in the GV-12m group due to the low number of detected CpG sites (0.025M). Then we calculated the mean methylation level of different types of cytosines covered with more than three reads. Surprisingly, the pattern of CpG sites and non-CpG sites were entirely different. While non-CpG sites gained methylation, demethylation of CpG sites was observed during GV oocyte aging (Figure 6a). The expression of non-CpG methylation-related enzyme, *Dnmt3a* was consistently increased in GV oocytes, which was consistent with the accumulation of methylation in non-CpG sites during GV oocyte aging (Figure 6b). In contrast, the age-related pattern of DNA methylation in MII stage was reversed. Global methylation level of CpG sites in MII oocytes increased with aging, but reduction of methylation was observed at non-CpG sites (Figure 6a). Our finding is consistent with the published result of the decline in global methylation level during MII oocyte aging measured by immunofluorescence [33]. During GV oocyte aging, the expression level of demethylase Tet1, was downregulated but Tet3 was up-regulated during MII oocytes aging (Figure 6b). Thus, age-related global methylation changes were consistent with the expression dynamics of demethylases, *Tet1* and *Tet3*, and methylases, *Dnmt3a*.

**Figure 6.**
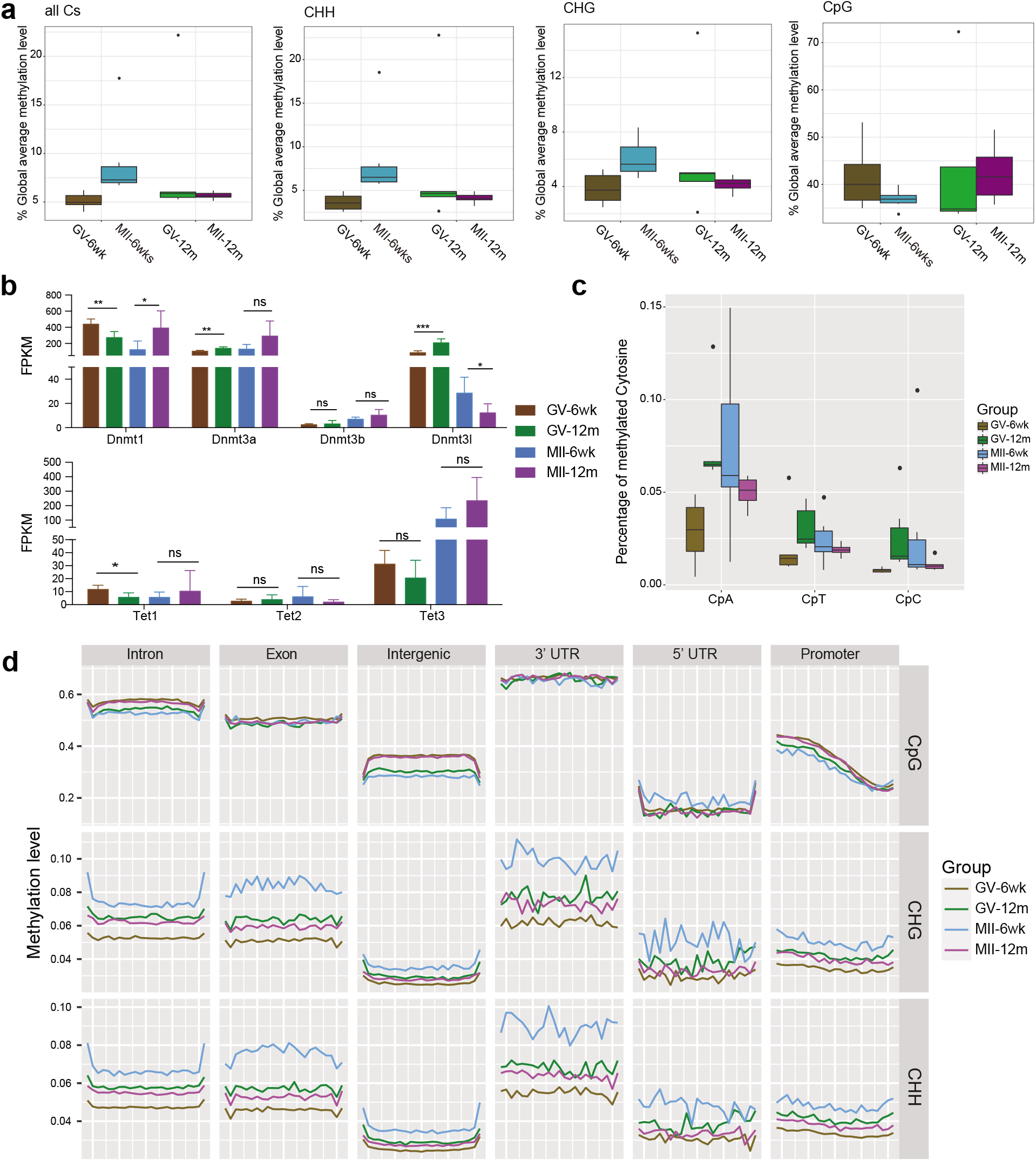
Global DNA methylation was influenced during oocyte maturation and aging. **(a)** Box plots of the global methylation of all sites, CHH sites, CHG sites and CpG sites (from left to right) in four groups (GV-6wk: brown; GV-12m: green; MII-6wk: blue; MII-12m: purple). The oocyte in GV-12m group with the lowest mapping efficiency is excluded. **(b)** Bar plots showing the expression of methylation associated genes in two groups. Comparisons are performed by One-way ANOVA (ns: not significant; *p<0.05; **p<0.01). **(c)** Box plot demonstrating the percentage of methylated cytosines in different di-nucleotide sequences (CpA, CpT and CpG) in four groups. **(d)** Composite plot of methylation levels across the genomic feature. The cytosine sites covered with less than three reads in the in silico merged group data have been filtered. The bin size of 200bp is applied to calculate the methylation level. Promoters are identified as the region of 2000bp upstream to TSS. The other genomic features are obtained from the UCSC genome browser. Different genomic contexts of cytosine, including CpG (Upper panel), CHG (Middle panel) and CHH (Low panel) are examined separately.

Moreover, the distinct pattern of maturation-associated DNA methylation changes was also demonstrated (Figure 6a). Global methylation level of non-CpG sites was accumulated during maturation in 6-weeks old mice, but declined in 12-months old mice. The maturation-associated changes of non-CpG sites in 6 weeks old mice was coherent with the published study of human oocytes [34]. Besides, the decline of CpG methylation during oocyte maturation was observed at six weeks old, but the opposite trend was found in 12 months old, which aligned with mRNA changes of a maintenance CpG methyltransferase, *Dnmt1*, during young and old oocyte maturation (Figure 6b).

On the other hand, we used the merged data to investigate whether any particular genomic feature had different patterns in comparison to the global pattern. The mean methylation levels of CpG, CHH and CHG sites were plotted according to different genomic regions and different groups (Figure 6d). The age-related or maturation-related methylation changes of CpG or non-CpG were consistent with the global pattern. It suggested that age-related or maturation-related dynamics of CpG and non-CpG methylation was unbiased to gene regions. We also detected the methylation in CpA was more common than the other non-CpG sites (Figure 6c), which was similar to a previous report [35]. Therefore, maternal aging could distort the normal global DNA methylation dynamics during oocyte maturation, and one potential mechanism was the changed expression of DNA methyltransferases caused by aging.

Next, we focused on the age effect on CpG methylation. A total of 1170 age-related DMRs were identified in GV oocytes. Among them, 385 DMRs gained methylation during GV oocyte aging (Hyper-CpG-DMR), while 785 DMRs were demethylated (Hypo-CpG-DMR). Similarly, during MII oocyte aging, 146 DMRs gained methylation and 79 DMRs lost methylation. The larger number of identified DMRs during GV oocyte aging implies GV oocytes are more susceptible to methylation changes during maternal aging. The majority of these DMRs were located in introns and intergenic regions (Additional file 1: Figure S4a). These DMRs were mapped to the nearest genes with the maximum region of 100kb upstream and downstream. Genes with age-related DMRs in GV oocytes showed functional enrichment in cell adhesion, while genes with age-related DMRs in MII oocytes were enriched in neuron differentiation (Additional file 1: Figure S4b). Unfortunately, we did not find any DMR with the same age-related change trend shared in the two ages.

Motif finding analysis on these age-related CpG DMRs in GV and MII stages revealed that these regions are enriched in binding sites of multiple shared transcription factors, with YY1 being the most significant one (Additional file 1: Figure S4c-d). YY1 is a component of ribonucleoprotein complexes and plays a role in the storage or metabolism of maternal transcripts [36]. A YY1 binding site in the promoter of *Spin2d* (Spindlin family member 2A), which is involved in cell cycle regulation and exhibited H3K4me3-binding activity, gained methylation during GV oocyte aging. Its family member, *Spin1d*, was essential for MII stage maintenance and chromosomal stability [37]. This implied that age-related methylation alteration in the YY1 binding site of Spin2d might affect H3K4me activation in GV oocytes. Moreover, several meiosis-related genes (*Ccng1* and *Mei4*) and a mitochondrion-associated gene (*Ndufb8*) demonstrated demethylation in their YY1-binding sites (Additional file 1: Figure S4e), which supported our hypothesis that maternal age might impair mitochondrial function and meiosis progress through DNA methylation alteration.

Besides CpG, the global pattern of non-CpG methylation was also altered during oocyte aging. We identified 2633 age-related hypermethylation DMRs (Hyper-nonCpG-DMR) in GV oocytes, and 2023 DMRs lost methylation (Hypo-nonCpG-DMR) for non-CpG. Meanwhile, in MII oocytes, 609 Hyper-nonCpG-DMRs and 1275 Hypo-nonCpG-DMRs were identified. The lower number of non-CpG-DMRs during MII oocyte aging confirmed the greater impact of age on DNA methylation in GV oocytes than MII oocytes. Similar to CpG-DMRs, the majority of non-CpG DMRs were annotated to intergenic genome and introns (Additional file 1: Figure S4f). Genes related to these nonCpG-DMRs showed prominent functions in neuron development and cell adhesion (Additional file 1: Figure S4g). When comparing age-related nonCpG-DMRs in two stages, we detected 18 common Hyper-nonCpG-DMRs and 12 common Hypo-nonCpG-DMRs. *Dlg2* (Discs large MAGUK scaffold protein 2) and *Msr1* (Macrophage scavenger receptor one) were involved (Additional file 1: Figure S4h), and their methylation was reported to be altered by aging or environment [38, 39].

### The interplay between transcriptome and methylome during oocyte aging

After studying the age effect on transcriptome or methylome in mouse oocytes respectively, we asked whether these methylation changes were associated with changes in gene expression. The Circos plots illustrated genome-wide changes of gene expression and DNA methylation during GV and MII oocyte aging (Figure 7a). In each oocyte group, we divided all detected genes into three expression classification based on their mean FPKM and plotted CpG methylation across genes. Globally, a positive correlation between CpG methylation in gene bodies and gene expression was shown (Figure 7b), which was consistent with the published study on other cells [40]. Next, we examined the interplay between gene expression and DNA methylation in individual genes. We calculated the mean CpG methylation level of the promoter or gene body for each gene in individual cells. Pearson’s correlation analysis on gene expression and DNA methylation was performed for each gene. We identified 138 genes demonstrating strong correlation between gene expression and methylation in promoters (Figure 7c). Among the 138 genes, 99 genes showed a positive correlation, and the other 39 demonstrated a negative correlation. The expression of *Stat3* (Signal transducer and activator of transcription 3), which activates the transcription of oocyte-specific genes (*Gdf9, Bmp15* and *Jagged1*) [41], was increased with the methylation level of its promoter (Pearson’s correlation coefficient = 0.9144, Figure 7d). Higher methylation level in the promoter of *Tarbp2* (Trans-activation responsive RNA-binding protein 2), which is a critical cofactor of Dicer and supports the maturation of highly expressed miRNAs in oocytes [42], resulted in lower expression level (Pearson’s correlation coefficient = −0.9976, p<0.05, Figure 7d). Meanwhile, we also detected 31 genes with positive correlation between gene expression and methylation level in gene bodies and 14 genes with negative correlation (Figure 7c). An example of the negative correlation was *Rxrb* (Retinoid X receptor beta), which is involved in retinoic acid (RA) signaling and RA-dependent chromatin alterations in Xenopus oocytes (Pearson’s correlation coefficient = −0.8861, Figure 7e) [43]. In contrast, a potential marker in the cumulus-oocyte-complex for predicting embryo morphology, *Stc2* (Stanniocalcin 2), showed a positive correlation between gene expression and methylation level in the gene body (Pearson’s correlation coefficient = 0.9550, Figure 7e) [44]. Above all, scM&T-seq not only allowed us to investigate the relationship between gene expression and methylation in cells, but also demonstrated a novel and gene-specific association across cells, which might challenge the conventional knowledge about the association between gene expression and DNA methylation established mainly on bulk samples.

**Figure 7.**
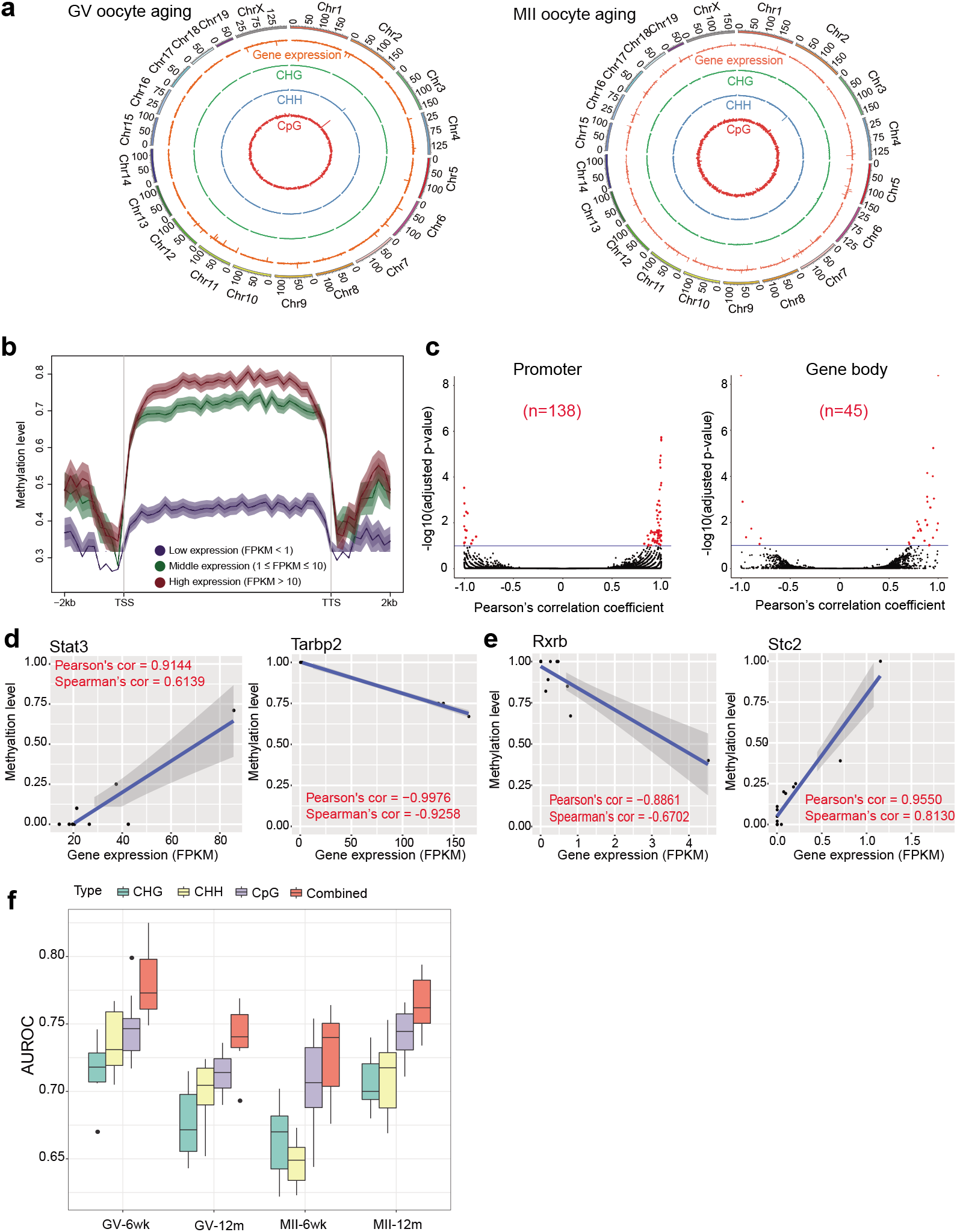
The interplay of DNA methylation and gene expression during oocyte aging. **(a)** Circos plots illustrating genome-wide changes of DNA methylation and gene expression during GV (Left) and MII (Right) oocyte aging. Chromosome ideograms are shown in the outermost ring of the circus plots. The inner orange rings represent age-related gene expression changes. The inner green, blue and red rings represent age-related methylation changes in CHG, CHH and CpG sites respectively. The scale of methylation changes is from −1 to 1. **(b)** CpG methylation level of genes with different expression across genomic regions in GV-12m group. Genes are divided into three groups based on FPKM. CpG sites covered with more than three reads in the pooled group data are involved. The methylation level is calculated in each 200bp bin across the region of 2000bp upstream to TSS and 2000bp downstream to TTS. Each line consists of the mean methylation with 95% confidence interval. **(c)** Volcano plots demonstrating the genes with the strong correlation between methylation level in promoters (Left) or gene bodies (Right). The x-axis in the volcano plots represents the Pearson’s correlation coefficient, and the y-axis represents p values adjusted by Benjamini and Hochberg (BH) method. Each dot signifies each gene and red dots show genes with significant correlation. **(d)** Scatter plots showing examples of genes with the strong correlation between methylation level in promoter and gene expression. **(e)** Scatter plots showing examples of genes with the strong correlation between methylation level in gene body and gene expression. Dots in (d) and (e) represent individual samples. The regression line is shown as blue and the 95% confidence interval is demonstrated in the grey region. **(f)** Cross-validation AUROC results of predicting gene expression class by cross-chromosome using gene body features.

Previous studies suggested gene body methylation is an indicator of gene expression levels in many types of cells. Thus, we investigated whether we could apply the methylation of gene body to predict the expression level of the corresponding gene in mouse oocytes. We divided each transcript into six regions, including first exon, first intron, internal exons, internal introns, last exon, and last intron, and computed the average methylation level of each region across all four groups (Ref. PMID 25074712). Expression of genes was classified to high or low using the median expression value as threshold. A Random Forest model was built to check whether the methylation levels at the six regions of a gene body could infer the expression class of the corresponding gene (Ref: Breiman L. Random forests[J]. Machine learning, 2001, 45(1): 5-32). By a rigorous cross-validation procedure, we saw that all the areas under the receiver operating characteristics (AUROC) across all the oocyte groups were significantly higher than the random background value of 0.5 (Figure 7f), indicating that the methylation levels of gene body could infer high or low expression classes globally. Moreover, among methylations of all cytosine in di-/tri-nucleotides, CpG served as a better indicator for gene expression classes than CHH and CHG did across all the oocyte groups, while methylation of all Cs together gave the best accuracy (Figure 7f, indicating that methylation of different cytosine di-/tri-nucleotides may contain non-redundant information about gene expression. We also tried building gene expression prediction models using promoter methylation levels as input but the results were not satisfactory (results not shown), suggesting that gene-body methylation is a better and more reliable indicator of gene expression in mouse oocytes.

### Dynamics of transposable elements (TEs) during oocyte aging

Transposable elements are regions containing repetitive sequences in the genome and were previously considered as “junk”. But recently, TEs have been implicated in the loss of genome integrity and gene regulation [45]. Therefore, we measured the expression levels and DNA methylation of TEs to investigate the effect of maternal aging on TEs. In both GV and MII oocytes, TEs showed higher correlation coefficient between two different ages in comparison to coding genes and all detected transcripts (Additional file 1: Figure S5a-b), suggesting that maternal age had a relatively minor effect on TEs expression in oocytes. Our data showed that TEs lost methylation during oocyte maturation, and maternal age retarded the maturation-related demethylation in TEs (Additional file 1: Figure S5c). We detected 183 TEs with differential expression during GV oocyte aging and 35 TEs during MII oocyte aging (Additional file 1: Figure S5d-f), which suggested that maternal age would exert more effect on the activity of TEs in GV oocytes than MII oocytes. Based on the classification of these differentially expressed REs, most of them were assigned to long terminal repeats (LTRs) and long interspersed nuclear element (LINEs, Additional file 1: Figure S5e,g), which indicated that LINEs and LTRs were the most susceptible to maternal aging. More interestingly, Gene ontology analysis on age- or maturation-related CpG DMRs demonstrated that LINE1 regions, especially the youngest subfamilies, L1Md_A and L1Md_T, were highly enriched (Additional file 1: Figure S5h). This suggested that maternal age could affect both transcription and DNA methylation in LINE1, which might contribute to the decline of oocyte quality in aged female. To the best of our knowledge, we are the first to pursue the changes of gene expression and DNA methylation in TEs during oocyte aging.

### Establishing a prediction model for mouse oocyte quality

We built a two-layer machine learning model incorporating gene expression levels to predict oocyte aging and maturation groups respectively(Figure 8a-b, Additional file 1: Figure S8a), with the first-layer learnt a small number of genes among the large set of predictors and the second-layer learnt the final scoring model. The first layer learnt important genes in predicting oocyte groups by Logistic Regression (LR) classifier with L1 norm penalty. In the first layer model, we achieved an average AUROC of 0.81 and 1.0 by repeating the whole cross-validation procedure 10 times for age group and stage group respectively (Figure 8c, Methods). The second layer used the top 10 critical genes of non-zero coefficients selected from the first layer and built another LR model using these genes. The learnt contributing importance scores of different genes were shown in Figure 8d-e.

**Figure 8.**
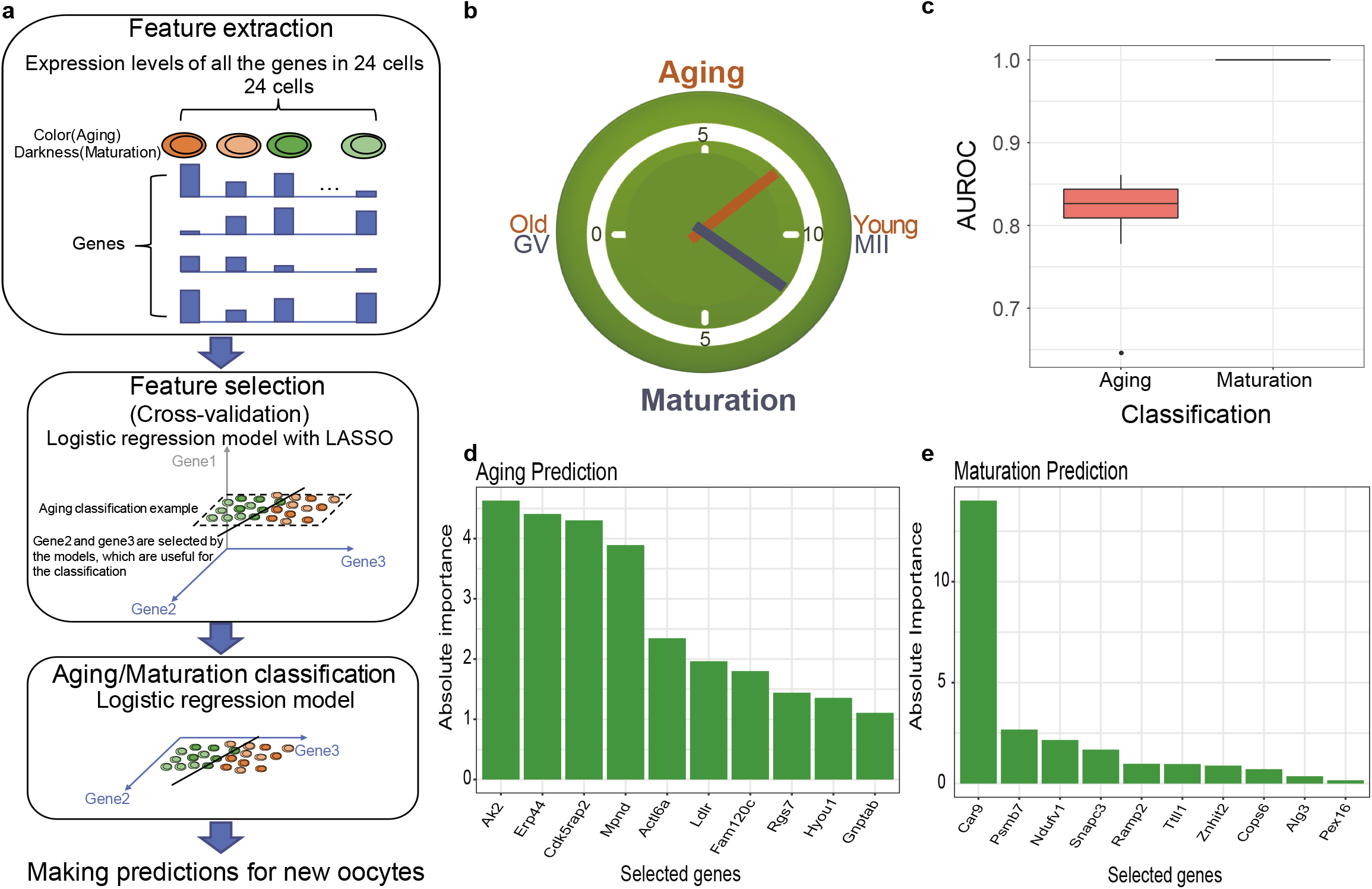
Establishment of the prediction model of oocyte quality. **(a)** A two-layer machine learning model incorporating gene expression levels to predict oocyte aging/maturation classes. **(b)** Oocyte aging/maturation clock. **(c)** AUROCs of cross-validation repeated ten times in feature selection. **(d-e)** Top 10 genes selected by our model ordered based on feature importance of aging prediction(f) and maturation prediction(g) respectively.

In the gene list of the aging prediction model, mitochondria-related genes, such as *Ak2* (Adenylate kinase 2), and meiosis-associated genes, such as *Cdk5rap2* (Cyclin-dependent kinase 5 regulatory subunit associated protein) and *Actl6a* (Actin like 6A), were included, which was consistent with the defects in mitochondria and chromosome segregation during oocyte aging observed in our scRNA-seq analysis (Additional file 1: Figure S6). *Hyou1* (Hypoxia up-regulated 1), which was downregulated during oocyte aging in our model (Additional file 1: Figure S7), was reported to be decreased in aged skin (http://www.ageing-map.org/). Meanwhile, several known maturation-associated genes, like *Ndufv1* (NADH ubiquinone oxidoreductase core subunit v1), *Psmb7* (proteasome subunit beta 7) and *Ramp2* (receptor activity modifying protein 2), were involved in the maturation prediction model.

Practically, given a new oocyte cell with the expression levels of at most 20 genes measured, our two models could predict which classes (young vs. old and GV vs. MII.) it belongs to by giving a confidence score ranging from 0 to 10. To test the feasibility, we downloaded a published mouse oocyte scRNA-seq data set (GEO: GSE70605) [46]. According to our prediction model, these oocytes were assigned to young and MII classification, which was consistent with the described sample information, that MII oocytes were collected from 10 weeks old mice (Additional file 2: Table S3). This indicated the feasibility and accuracy of our prediction models.

## Discussion

We are the first to employ the single cell parallel sequencing approach to simultaneously profile aging-associated transcriptome and methylome from single oocytes. In this study, we revealed how maternal aging affected various cellular mechanisms in oocytes at different stages, which filled the knowledge gap of the molecular dynamics during oocyte aging. We found that mitochondrial dysfunction played a critical role during GV oocyte aging. Previously, several studies demonstrated that maternal aging affected mitochondrial quantity and quality in oocytes. ATP, the major product of mitochondria, was reported to be significantly decreased in mouse aged GV oocytes, though mtDNA copy number was not changed significantly during GV oocyte aging [47]. It implied that the reduction in mitochondrial activity during GV oocyte aging was resulted from not only the decreased number of mitochondria but also poorer mitochondrial quality. Our study provided molecular evidence of disordered mitochondrial activity in aged GV oocytes. This suggested that the recruitment of mitochondrial dysfunction in aged GV oocytes would be a promising therapeutic target for poor oocyte quality in aged women. Alternatively, our study demonstrated that the deterioration in spindles and microtubules were prevalent in aged MII oocytes. Chromosome segregation is a prominent step in the maturation to MII oocytes. The formation of spindle and microtubule is essential to maintain equal segregation of chromosome during cell division, and microtubule pushing has been recently proved to be indispensable to chromosome separation [48, 49]. Higher frequency of aneuploidy was also widely observed in aged MII oocytes [50]. Our data suggested that abnormal expression of microtubule-associated genes might contribute to dysfunction in chromosome separation during MII oocyte aging. Besides microtubules, spindle assembly is also an essential step during meiosis. DEGs during MII oocyte aging were enriched in spindle assembly and the reduced expression of spindle assembly checkpoint gene, *Mad2*, was observed in our data, which was consistent with published human bulk sample data [51]. Thus, our findings brought out different research directions to further study maternal aging associated molecular mechanisms in oocytes at different stages.

Besides changes at a particular stage, maturation-associated transcript dynamics has not received much attention in studies related to oocyte maternal aging. In this study, we demonstrated that the maturation-associated alterations in transcriptome were affected by maternal age. Among them, EIF2 protein synthesis pathway was the most vulnerable. This implies that defects in protein synthesis occurred in the oocyte maturation transition during maternal aging, thus protein dynamics should play a role in aging-dependent decline of oocyte quality. Schwarzer et al. carried out the first study of proteome dynamics in mouse oocytes during maternal aging and concluded that proteome analysis could not be surrogated with transcriptome analysis due to the significant difference between them [52]. Reyes et al. had recently compared gene expression changes of maturation between young and advanced maternal age human oocytes and showed DEGs were enriched in oxidative phosphorylation, mitochondrial dysfunction and DNA replication [53]. However, during DEG identification in their study, genes with huge expression changes in different ages were not included, and genes with reversed expression changes in different ages were not distinguished from functional analysis. These disadvantages limited the accuracy of their conclusion. To overcome the above shortcomings, we established a linear logistic regression model to identify genes which expression pattern during maturation was discrepant in young and aged oocytes. Our approach is expected to provide more unbiased and dependable conclusions.

In addition, we identified IL-7 as a potential molecular target to improve aged oocyte maturation. IL-7 is an oocyte-secreted factor during oocyte maturation. Upon stimulation of FSH, cumulus cells enhance the secretion of IL-7 of their attached oocyte through promoting its mRNA translation. The secreted IL-7 in follicular fluid binds to the IL-7 receptors on the surface of cumulus cells and then induce cumulus cell proliferation [Ref PMID 26864200]. Previous studies showed that the level of IL-7 in the polysome fraction increased during oocyte maturation, and IL-7 protein content increased in the follicular fluid during oocyte maturation. They also demonstrated that the IL-7 content in the follicular fluid of normally fertilized eggs was higher than that of abnormally fertilized ones [54]. It was shown that accumulation of IL-7 protein in the medium of *in vitro* maturation is correlated with oocyte quality improvement [55–57]. In both our data and published microarray data, the expression level of IL-7 was decreased during oocyte aging, which suggested that the reduction of IL-7 in oocyte might contribute to the declined oocyte quality. Therefore we propose that supplement of IL-7 for *in vitro* maturation of aged oocytes might improve the maturation of aged oocytes and rescue their low development competency.

Besides aging-related dynamics in transcriptomes, we demonstrated the accumulated methylation in non-CpG sites during GV oocytes aging. Methylation accumulation in non-CpG sites was also observed in aging-related diseases such as type 2 diabetes [58]. *Dnmt3b* and *Dnmt3a* were believed to participate in methylation of non-CpG sites in ESCs and neuron respectively [59]. Our scRNA-seq data revealed the upregulation of *Dnmt3a* and *Dnmt3b* during GV oocyte aging, which implied the involvement of these two methyltransferases in non-CpG methylation regulation in GV oocytes. The expression levels of *Dnmt3a* and *Dnmt3b* was not altered during aging of MII oocytes according to our data (Figure 6b). Our data suggests the potential aging-related post-transcriptional regulation mechanisms on methyltransferases resulting in the defects in DNA methylation in MII oocytes. More importantly, these findings provide additional support for cell type-specific regulation of non-CpG methylation.

We also observed the hypermethylation of CpG sites with increased expression of *Dnmt1*, a methyltransferase responsible for CpG methylation maintenance, during MII oocytes aging (Figure 6b). A recent study demonstrated that Dnmt1 also functions as a de novo methyltransferase during oogenesis (Ref: https://www.nature.com/articles/s41586-018-0751-5). These results suggested that aberrant expression of *Dnmt1* induced by maternal age could contribute to exceeded methylation establishment in MII oocytes.

Other than studying the maternal aging effect on particular oocyte stages, we also demonstrated the global hypermethylation pattern during young oocyte maturation, which was consistent with the previous understanding [34]. However, this pattern was disrupted during aged oocyte maturation. Methylation maintenance by *Dnmt1* was disturbed according to its disordered expression pattern during aged oocyte maturation (Figure 6b). Thus, we propose that maternal aging would disrupt DNA methylation regulation during oocyte maturation through both *de novo* methylation and methylation maintenance mechanisms.

The simultaneous generation of the transcriptome and DNA methylome profiles from the same cell allowed us to explore the regulation of DNA methylation in transcriptome. We demonstrated a positive correlation between gene expression and methylation in gene body, but not in promoter in oocytes, which was in good agreement with previous study [62]. A possible explanation was due to the relatively high global methylation level in oocytes than other somatic cells. Besides, we also illustrated the relationship between methylation level in promoter or gene body and expression for individual genes in oocytes varies. Besides DNA methylation, other regulation mechanisms, like histone modification, chromatin accessibility and transcription factor binding are involved in determining gene expression [63, 64]. Effect of individual epigenetic mechanism on gene regulation is lower than expected, which could also be concluded from the previous study of single-cell multi-omics sequencing in ESCs [65, 66].

After identifying the dynamics in transcriptome and methylome during oocyte aging, we developed a prediction model for oocyte quality. Due to low genome coverage of DNA methylation data, a model based on oocyte transcriptome was established. Our model has excellent performance in both cross-validation and test set from public datasets. Moreover, the L1 norm term helps to identify important genes that contribute most in distinguishing oocytes of different ages and maturation stages based on feature importance. It could not only lead to potential interesting discoveries, but also economically guide the design of biological experiments. Compared with other computational and statistical models based on unsupervised schemes such as the p-values of the differentially expressed genes, the prediction score of our model quantifies the oocyte age or maturation stages into a 1-10 scale and can be interpreted easily. One of the future direction of our model is to extend the current binary classification task to multi-class classification or regression, such that it can predict, the exact age of any given oocyte, rather than whether it is young or old only. Finally, our model is an open and flexible framework that can be easily adjusted for human oocytes by replacing the gene expression data from mouse with those from human. We believe this could potentially serve to facilitate clinical decisions.

## Conclusions

In conclusion, we generated comprehensive profiles of simultaneous transcriptome and methylome in mouse oocytes of different stages and ages at single-cell and single-base resolution to decode aging-associated multi-omics alterations and regulatory mechanisms in oocytes contributing to oocyte quality. The discoveries in our study provide the promising candidate therapies for rescuing or improving quality of aged oocytes. Remarkably, to the best of our knowledge, we are the first to establish the oocyte quality prediction model based on gene expression, which could overcome the limitation in recent morphology assessment and will be translated to human oocytes for clinical application in the future.

## Materials and Methods

### Animals and oocyte collection

C57/BL6 female mice aged 6 weeks old (6wk) and 12 months old (12m) were used in this study. All the experimental procedures on animals were approved by the Animal Experimentation Ethics Committee of The Chinese University of Hong Kong. To collect GV oocytes, mice were injected intraperitoneally with 10 IUpregnant mare’s serum gonadotropin (PMSG, Sigma) 48 hours before sacrifice. Ovaries were dissected from sacrificed mice and mashed in a dish containing M2 medium (Sigma). GV oocytes attached to cumulus cells were transferred in droplets containing M2 and 10% (mass: volume) hyaluronidase (Sigma) for detachment. Then, cumulus cell-free GV oocytes were washed three times in M2 medium following in PBS three times. Each oocyte was finally transferred into individual PCR tubes with less than 1 μl of PBS by mouth pipette and stocked in −80°C freezer. For MII oocytes collection, mice underwent the full superovulation procedure with an intraperitoneal injection of 10 IU human chronic gonadotropin (hCG, Sigma) 48 hours after PMSG administration. 14-16 hours after hCG injection, female mice were decapitated to obtain tissues including ovary, oviduct and part of the uterus. These tissues were put in a dish containing M2 medium, and the ampulla of oviduct was chopped by a 29G needle. Cumulus-oocyte-complex (COC) were removed from the oviduct and transferred to M2 medium containing 10% hyaluronidase. The following wash steps and storage method for MII oocytes were the same as GV oocytes. In this study, GV oocytes and MII oocytes were collected from both 6 weeks old and 12 months old mice and were accordingly assigned to one of the four oocyte groups, namely GV-6wk, GV-12m, MII-6wk and MII-12m.

### Single cell M&T sequencing

The modified single-cell M&T-seq protocol was employed to generate simultaneous transcriptome and methylome profiles from single oocytes [67]. Briefly, 7 μl of soft lysis buffer containing 500 mM KCl (Sigma), 100mM pH 8.0 Tris-HCl (Sigma), 1.35 mM MgCl2 (Sigma), 4.5 mM DTT (Thermo Fisher Scientific), 0.45% Nonidet P-40 (Sigma), 0.18U SUPERase-In (Thermo Fisher) and 0.36U recombination RNase-inhibitor (Thermo Fisher) were added into each PCR tube containing a single oocyte and were incubated at 4°C for 30 min. Then, the lysate was vortexed at room temperature for 1 min and centrifuged at 1000g for 5 min to allow the nucleus to sink to the bottom of the tube. 6.5 μl of supernatant containing RNA was carefully transferred to a new PCR tube for generating scRNA-seq library, and the remaining 1.5 μl of lysis solution with the nucleus was subjected to scBS-seq.

To prepare scRNA-seq libraries, SMART-Seq2 was applied to cDNA preparation and pre-amplification from RNA. Five nanograms of amplified cDNA per sample quantified by Qubit 3.0 Fluorometer (Thermo Fisher) was used to construct RNA-seq libraries using TruePrep DNA Library Prep Kit V2 for Illumina (Vazyme). For preparing scBS-seq, 5 μl of RLT plus buffer (QIAGEN) and 60 fg of λ-DNA were added into each isolated nucleus to lyse nucleus completely. The subsequent bisulfite conversion, amplification and library construction procedures were performed with Pico Methyl-Seq^TM^ Library Prep Kit (Zymo Research) following the manufacturer’s instruction. Finally, the quality of scRNA-seq and scBS-seq libraries was checked by Bioanalyzer 2100 (Agilent) using High-sensitivity DNA chips, while the yield of libraries was measured by Qubit 3.0 Fluorometer. The qualified libraries were multiplexed and sequenced on Illumina HiSeq2500 platform for generating 150bp pair-end reads.

### scRNA-seq data analysis

Raw reads from both RNA-seq and scBS-seq were checked by FastQC (v0.11.8, https://www.bioinformatics.babraham.ac.uk/projects/fastqc/) and trimmed by Trim Galore (v0.4.5, https://www.bioinformatics.babraham.ac.uk/projects/trim_galore/). For RNA-seq data, trimmed reads were mapped to the mouse reference genome (mm10) by TopHat (v2.1.1) [68] with default parameters and Cufflinks was employed to quantify gene expression level with gene annotation obtained from Ensembl GRCm38.p5 [69]. Gene expression level was calculated in the form of fragments per kilobase million (FPKM). Genes with undetected expression level in all the samples were excluded in the following analysis.

#### Principal component analysis (PCA)

Gene expression matrix was generated from the output of Cufflinks in the form of normalized FPKM. All qualified genes were used for PCA with R function pcromp. Top three principal components (PC1, PC2 and PC3) were applied to plot the 3D figures by R.

#### Differential expression genes (DEGs) detection

Differential expression of genes among the four groups was calculated by Cufflinks with default parameters. Genes with fold change >2 or <0.5 and Benjamini and Hochberg (BH) adjusted p-value less than 0.05 were identified as differentially expressed genes (DEGs).

#### Function analysis on DEGs

Identified DEGs were used for Gene Ontology (GO) and Kyoto Encyclopedia of gene and genome (KEGG) pathway analysis with R package, clusterProfiler [70].

To further analyze functions and potential links of DEGs, Ingenuity Pathway Analysis software (IPA, QIAGEN Inc, www.qiagen.com/ingenuity) was applied for further analysis. The list containing both identified DEGs and their corresponding log2-transformed value of fold change was put into the software. Based on the Ingenuity Knowledge Base and built-in algorithms, canonical pathways enriched in DEGs were calculated and demonstrated. Besides, according to known gene-gene regulation or connectivity in the Ingenuity Knowledge Base and the actual gene expression dynamics in our data, several upstream regulators and their function were predicted and represented as network plots.

#### Isoform switch analysis

The expression level of transcripts was quantified by RSEM [71] and was used as the input of R package, IsoformSwitchAnalyzeR, to identify potential aging-related isoform switches [32]. Transcripts with the extremely low expression level of TPM (transcripts per million) <0.1 were filtered out in advance. Isoform Fractions (IF) was applied to describe the usage of one isoform in its parent gene. IF was compared in the pairs of groups to test the difference in isoform usage. Differences in isoform usage with adjusted p-value < 0.05 was considered as isoform switches. Identified switches were subjected to predict their biological function and consequences based on sequences. Besides, the biological mechanisms of isoform switch, including alternative splicing, alternative transcription start sites (TSS) and transcription termination sites (TTS), were predicted.

#### Repetitive elements expression

The expression level of repetitive elements was quantified by RSEM with the reference annotation file downloaded from RepeatMasker (v4.0.7, https://www.repeatmasker.org).

### scBS-seq data processing

Trimmed scBS-seq data was aligned to mm10 genome by Bismark (v0.20.0) by single-end mode to obtain higher mapping efficiency [72]. Two bam files obtained from one sample were merged together and the merged bam files were deduplicated and used to calculate methylation level following the manual of Bismark. Only the cytosine sites covered with more than 2 reads were included in the following analysis. Bisulfite conversion efficiency was calculated by the methylation level of unmethylated λ-DNA. Samples with bisulfite efficiency less than 95% were ruled out. To reduce amplification bias and sequencing bias, we assigned cytosine sites whose methylation level was equal to or less than 0.3 to be unmethylated. Similarly, cytosine sites with methylation level ≥0.7 were considered as methylated. Descriptive statistical analysis, such as read coverage per base and base annotation, were performed by R package methylKit [73]. When calculating the methylation level across the genomic features, mean methylation level of CpG, CHH and CHG sites were calculated in the bin size of 200bp in each group. SeqPlots was employed to perform the analysis and to draw plots [74].

To increase genome coverage, we combined all detected Cs in samples of the same groups. Differential methylation regions (DMRs) were identified using metilene [75]. Detected DMRs were annotated by the command annotatePeak.pl in HOMER (http://homer.ucsd.edu/homer/ngs/quantification.html) based on Ensemble GRCm38.p5. The enrichment analysis of DMRs was performed by R package, annotatr [76]. GO analysis for genes with DMRs were performed by clusterProfiler R package.

### Interplay between DNA methylation and gene expression

#### Correlation between gene expression and DNA methylation across genes

The expression level of each gene was calculated as the average of samples in the same group. CpG methylation level was merged from samples in the same group as described above. Methylation level of different groups of genes was calculated in bin size of 200bp. Genes were classified into three groups consisting Low expression group (FPKM ≤ 1), Median expression group (1 < FPKM < 10) and High expression group (FPKM ≥10). The plots were generated by SeqPlots.

#### Correlation between gene expression and DNA methylation across oocytes

The pairs of gene expression and its corresponding methylation level of different genomic regions (including promoter and gene body) in every single cell were listed as a matrix. Genes detected in less than four cells were discarded. Pearson’s correlation was performed to assess the correlation between gene expression and DNA methylation of different genomic regions for each gene, and P-values were adjusted for multiple testing by Benjamini-Hochberg method. The promoter was defined as the region from 2000bp upstream of TSS to 1000bp downstream of TSS. Gene body was considered as the region from TSS to TTS.

### Establishment of the oocyte quality prediction model

Logistic Regression (LR) model with L1 norm penalty (Formula 1) was built using gene expression data to predict the classes (6 weeks vs. 12 months and GV vs. MII) of the oocytes (Additional file 1: Figure S8). To predict age group, we grouped G6 and M6 as positive class and G12 and M12 as a negative class. A binary classifier was built using LR with L1 norm penalty. Among 24 oocytes, 23 were used each time for training and the one left for testing. To ensure a fair estimation of our model, we repeated the whole procedure for ten times, and thus in total, we had 24 × 10=240 models. The penalty coefficient λ was learnt during each of the cross-validation process based on 23 cells and λ that gave the minimum mean cross-validated error was applied in testing. ROC was computed to estimate model accuracy in each repeat (Figure 8b). We kept all the genes of non-zero coefficients in all the 240 models and counted how many times each gene occurred.

To estimate the probability distribution of how many times a gene would be selected randomly, we constructed a null model in the following way: In each of the 240 models for each classification task, we built a null model by selecting the same number of genes as the genes of non-zero coefficients selected by the true model. We repeated the whole procedure for ten times and plotted the probability distribution (Additional file 1: Figure S8b-c). At the p-value threshold 5e-4, in total, we obtained 75 significant genes for age classification and 44 significant genes for stage classification.

In terms of computational modeling, several top significant genes were selected in the majority of the 240 models, indicating that they are more general and useful for prediction. Moreover, for further potential biological validation experiments, we also picked the most promising genes in order to design the experiments effectively and economically. So we selected the top 10 genes among the significant genes which were most frequently selected by the LR model. Finally, we used the expression levels of the ten genes to predict oocyte age class by LR without any penalty term. Stage classes (GV vs. MII) were predicted in the same way. We also evaluated the contributing importance of the ten genes based on learnt coefficients (Figure 8 d-e). The whole procedure of the algorithm was implemented by R package Glmnet 2.0-16.

In addition, we tried to build a pure aging prediction model for 12 GV and 12 MII oocytes, respectively. Before feeding the genes to LR model, we first filtered out those genes which were lowly expressed and had lower variance (FPKM<1 and standard deviation < 1 in all 12 samples in GV or MII) since we only had half of the sample size (i.e. 12 samples) for each model, which may deal with fewer regressors. The average ROCs of GV and MII were 1 and 0.66 in the 10-fold cross-validation setting. The results indicated that the difference between the young and old oocytes in GV stage are large while the young and old oocytes in MII stage are similar, which was supported by a previous study [77]. It is also important to include larger size of oocyte samples for making confident predictions.

Formula 1 Objective function of Logistic Regression with L1 norm penalty

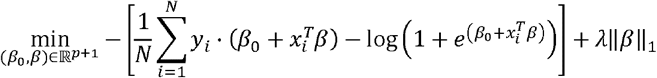

### Gene expression analysis by quantitative real-time PCR

The remaining cDNA after scRNA-seq libraries construction were subjected to quantitative real-time PCR. RT-PCR was performed using Power SYBR Green PCR Master Mix (Life Technologies, USA) on ABI ViiA-7^TM^ real-time PCR system with the standard relative quantification mode (2^ΔΔCT^) followed the instructions. The PCR system was initiated by heating 95°C for 10 minutes, followed by 40 cycles of 15 seconds at 95°C and 60 seconds at 60°C. The relative gene expression level was normalized by Gapdh. For each sample, three technical replicates were carried out on the same plate, with 3-4 biological replicates in each group. The sequences of PCR primers used were listed in Additional file 2: Table S4

### Statistical analysis

The expression data were shown as (mean ± standard deviation) unless otherwise mentioned. The comparison of expression results which were not mentioned in the above relevant section applied One-Way ANOVA among the four groups and was conducted by Prism 6. The definition of significance was following: *:<0.05; **:<0.01.

## Declarations

### Ethics approval and consent to participate

The study involving live mice was performed in accordance with the recommendations and guidelines of the CUHK Animal Experimentation Ethics Committee (AEEC), with all protocols approved by the Hong Kong Government Department of Health (Reference number: 17-459 in DH/SHS/8/2/1 Pt.3).

### Consent for publication

Not applicable

### Availability of data and material

The datasets generated and analyzed during the current study are available in the NCBI’s Gene Expression Omnibus repository (GEO119906, https://www.ncbi.nlm.nih.gov/geo/query/acc.cgi?acc=GSE119906).

### Competing interests

The authors declare that they have no competing interests.

### Funding

This work was supported by the Lo Kwee-Seong Biomedical Research Fund, School of Biomedical Sciences, CUHK and the Collaborative Research Fund (C4054-16G), Research Grants Council.

### Authors’ contributions

YQ designed the study, performed experiment, analyzed and interpreted data, and prepared the manuscript. QC was mainly responsible for sequencing data analysis and the establishment of oocyte quality prediction model and participated in manuscript writing. JL, AS and AL helped for animal keeping and manuscript preparation. NT, HC, KC, TCL, CW, CD, TLL, and KY provided intellectual inputs and advised on data interpretation. TLL was responsible for conceptual design and study coordination, interpreted data, and prepared the manuscript. All authors read and approved the final manuscript.

## Acknowledgements

The authors acknowledge the support of Core laboratory in School of Biomedical Sciences, The Chinese University of Hong Kong.

## Notes

### Competing Interest Statement

The authors have declared no competing interest.

