## Supplementary figures legend for "Simultaneous transcriptome and methylome profiles of single mouse oocytes provide novel insights on maturation and aging"

**Additional file 3: Supplementary figures legend**

**Figure S1 Correlation coefficients among samples and validation of scRNA-seq results by real-time PCR**

Heatmap plot showed correlation coefficients among all samples analyzed by Pearsons’ correlation (a). scRNA-seq results of randomly picked genes in single GV (b) or MII (c) oocytes from mice at 6 weeks and 12 months. (d) and (e) demonstrate the relative expression level of genes measured by real-time PCR in the corresponding cDNA left during library construction. (Mann-Whitney *U* test, *: p <0.05; **: p<0.01).

**Figure S2 Identification of TNF and PMA as upstream factors of aging-associated DEGs in GV oocytes by IPA.**

IPA prediction of upstream molecules, tumor necrosis factor (TNF, a) and phorbol myristate acetate (PMA, b) for aging-associated DEGs in GV oocytes is displayed. Legend key of IPA prediction is showed in the right.

**Figure S3 Identification of KDM5B as upstream factors of aging-associated DEGs in MII oocytes by IPA**

IPA prediction of an upstream molecule, KDM5B, for aging-associated DEGs in MII oocytes is demonstrated. Legend key of IPA prediction is showed in the right.

**Figure S4 Identification and characterization of age-related differentially methylated regions**

**(a)** Column diagrams demonstrating the annotation of identified age-related CpG-DMRs in GV or MII stages to different genomic regions. Age-related DMRs were divided into Hyper- or Hypo- based on their methylation changes. The analysis was performed by annotater. **(b)** Column diagrams showing GO analysis on genes nearest to identified CpG-DMRs. The x-axis represents the significance of enrichments in -log10(p-value). The top 8 enriched GO terms (all if less than 8 were identified) for each set of DMRs are shown. **(c-d)** Motif analysis of age-related DMRs in GV (C) or MII (D) stages. The sequences, names and corresponding p values of top 5 enriched motifs are listed. (e) Gene tracks showing the gain of methylation in the YY-1 domain of Mei4 during GV oocyte aging. The top two tracks showed the methylation level of CpG sites in GV-6wk and GV-12m group. The red line meant the full methylation, the blue one indicated non-methylation and the green one showed the half-methylation. The middle two tracks showed the peaks of YY1 Chip-seq data obtained from GSM2645432 and GSM 1665555). The bottom track demonstrated the gene structure of *Mei4*. **(e-f)** Column diagrams demonstrating the annotation of identified age-related nonCpG-DMRs in GV or MII stages to different genomic regions. **(g)** Example of common age-related Hyper-nonCpG-DMR locus in *Dlg2* (Left) and *Msr1* (Right). The methylation profiles of nonCpG sites are shown as heat maps. Dark blue represents sites of full methylation and white represents unmethylated sites. DMR loci are highlighted in red frames.

**Figure S5 LINE1 having the age-related susceptibility in transcription and DNA methylation**

**(a-b)** Scatter plots demonstrating expression level of genes and transposable elements (TEs) during GV (a) and MII (b) oocyte aging. Each black and red dot represents an individual gene and TE respectively. Pearson’s correlation between expressions at 6 weeks and 12 months were calculated for genes, TEs, and all transcripts. . **(c)** Bar chart showing the CpG methylation level of different TE families in four oocyte groups. **(d)** and **(f)** Volcano plots illustrating age-related differentially expressed TEs in GV (d) and MII (f) stage. Red dots represent TEs with significantly different expression during aging. **(e)** & **(g)** Column diagrams showing the classification of identified TEs with significantly different expression during GV (e) or MII (g) oocyte aging. **(h)** Dot plots representing Genome ontology analysis on identified age- and maturation- related differential expression TEs. The color of dots shows the enrichment scores, log_2_(observed/expected). The x axis shows the significance of the enrichment as –log10(p-value). The top 6 enriched genomic regions (all if less than 6 were identified) are listed for each set of TEs.

**Figure S6 Expression of genes selected in the prediction model of oocyte aging**

Box plots showing expression level (FPKM) of 10 genes involved in the prediction model of oocyte aging. All the oocyte samples from the same age (6-weeks-old, red & 12-months-old, green) were grouped together in this calculation.

**Figure S7 Expression of genes selected in the prediction model of oocyte maturation**

Box plots showing expression level (FPKM) of 10 genes involved in the prediction model of oocyte maturation. All the oocyte samples from the same stage (GV stage, red & MII stage, green) were grouped together in this calculation.

**Figure S8 Workflow of establishing the prediction model of oocyte aging and maturation.**

(a) Detailed workflow of establishing the prediction model of oocyte aging and maturation. **(b-c)** The probability of the times a gene is selected based on null models of aging prediction(b) and maturation prediction(c).
