## Supplementary material for "Simultaneous transcriptome and methylome profiles of single mouse oocytes provide novel insights on maturation and aging": Table S1-S4

**Additional file 2: Table S1-S4**

**Table S1 Performance of scRNA-seq on individual oocytes by single-cell M&T-seq**

| Group | Sample ID | Raw reads | Mapping rate (%) | Multiple alignment rate (%) | No. of detected genes |
| --- | --- | --- | --- | --- | --- |
| GV-6wk | R-6-G-1 | 6441156 | 64.1 | 7.8 | 20535 |
|  | R-6-G-2 | 6037417 | 65 | 12.9 | 20584 |
|  | R-6-G-3 | 9876427 | 62.9 | 12.2 | 22329 |
|  | R-6-G-4 | 7257003 | 64.4 | 10 | 20755 |
|  | R-6-G-7 | 6964803 | 62.5 | 10.1 | 20252 |
|  | R-6-G-8 | 6231430 | 62.7 | 8.8 | 21496 |
| MII-6wk | R-6-M-1 | 8490298 | 66.1 | 3.7 | 17112 |
|  | R-6-M-2 | 10842758 | 64.1 | 2.2 | 18860 |
|  | R-6-M-4 | 7421185 | 43.5 | 2.2 | 16617 |
|  | R-6-M-5 | 6038967 | 64.4 | 2.4 | 16633 |
|  | R-6-M-6 | 7007558 | 57.8 | 4.3 | 15260 |
|  | R-6-M-7 | 7934306 | 66.9 | 2.4 | 18096 |
| GV-12m | R-12-G-1 | 7950276 | 63.8 | 9 | 21188 |
|  | R-12-G-3 | 11501581 | 60.4 | 12.8 | 21024 |
|  | R-12-G-4 | 7240594 | 65.5 | 14.9 | 20363 |
|  | R-12-G-5 | 8925392 | 64.4 | 10.7 | 20824 |
|  | R-12-G-6 | 13321123 | 60.3 | 13.8 | 23890 |
|  | R-12-G-8 | 487223 | 58.8 | 13.8 | 12073 |
| MII-12m | R-12-M-1 | 8741826 | 56.9 | 2.3 | 18928 |
|  | R-12-M-2 | 5102691 | 47.5 | 2.4 | 15474 |
|  | R-12-M-3 | 4447166 | 60.5 | 2.5 | 15764 |
|  | R-12-M-4 | 3588340 | 34.3 | 4.4 | 12206 |
|  | R-12-M-5 | 3650323 | 31.6 | 4.9 | 12555 |
|  | R-12-M-7 | 7484372 | 26.4 | 5.8 | 14059 |

**Table S2 Performance of scBS-seq on individual oocytes by single-cell M&T-seq**

| Group | Sample ID | Total raw reads | Mapping efficiency (%) | Duplication rate (%) | Bisulfite conversion efficiency (%) | No. of CpG sites | No. of CHH sites | NO. of CHG sites |
| --- | --- | --- | --- | --- | --- | --- | --- | --- |
| GV-6wk | D-6-G-1 | 33583605 | 46.04 | 81.54 | 98.73 | 2532416 | 37302114 | 11161089 |
|  | D-6-G-2 | 35268606 | 42.18 | 87.01 | 98.77 | 1565663 | 24227014 | 6873537 |
|  | D-6-G-3 | 57195669 | 43.71 | 84.58 | 98.52 | 3114382 | 48339390 | 13864942 |
|  | D-6-G-4 | 38556854 | 54.50 | 79.84 | 98.76 | 3889270 | 56681007 | 17222183 |
|  | D-6-G-7 | 27515858 | 44.99 | 84.74 | 98.73 | 1531632 | 23295804 | 6723003 |
|  | D-6-G-8 | 34777937 | 47.14 | 84.17 | 98.58 | 2038975 | 32066760 | 9113342 |
| MII-6wk | D-6-M-1 | 28406376 | 52.22 | 97.6 | 98.76 | 195238 | 3346726 | 821505 |
|  | D-6-M-2 | 40585208 | 39.34 | 96.73 | 98.52 | 334168 | 5425439 | 1448320 |
|  | D-6-M-4 | 45865212 | 46.35 | 96.52 | 98.50 | 466563 | 7567480 | 2014111 |
|  | D-6-M-5 | 36230616 | 4.21 | 71.29 | 98.63 | 376219 | 6239682 | 1681687 |
|  | D-6-M-6 | 31469292 | 43.81 | 95.98 | 97.12 | 336158 | 5640606 | 1450350 |
|  | D-6-M-7 | 34430083 | 33.80 | 99.11 | 99.06 | 49294 | 1478280 | 180446 |
| GV-12m | D-12-G-1 | 23359356 | 40.11 | 86.25 | 98.80 | 1087092 | 15554011 | 4695049 |
|  | D-12-G-3 | 16872888 | 39.33 | 84.43 | 98.81 | 844375 | 12399672 | 3647340 |
|  | D-12-G-4 | 33306429 | 6.70 | 96.73 | 98.67 | 24654 | 577255 | 94798 |
|  | D-12-G-5 | 46849294 | 34.82 | 89.22 | 98.60 | 1296589 | 19843062 | 5653328 |
|  | D-12-G-6 | 32548430 | 25.00 | 98.87 | 98.83 | 64438 | 918374 | 234209 |
|  | D-12-G-8 | 13890717 | 32.45 | 86.67 | 97.96 | 488698 | 7022397 | 2144225 |
| MII-12m | D-12-M-1 | 35542929 | 38.25 | 84.2 | 98.55 | 1814540 | 26288374 | 7947412 |
|  | D-12-M-2 | 22034824 | 38.18 | 71.49 | 98.31 | 2237442 | 31175270 | 9623336 |
|  | D-12-M-3 | 27235723 | 43.97 | 84.18 | 98.28 | 1605774 | 22890927 | 6933733 |
|  | D-12-M-4 | 34146175 | 39.14 | 87.3 | 98.68 | 1411021 | 20248509 | 6121679 |
|  | D-12-M-5 | 3776127 | 18.34 | 56.64 | 98.39 | 451083 | 5907223 | 1910903 |
|  | D-12-M-7 | 3511491 | 31.62 | 75.87 | 98.64 | 228398 | 3262418 | 989367 |

**Table S3 Comparison between predict results and actual condition of oocytes in published data**

| Published Mouse Oocyte | Actual Age | Predicted Age  (score) | Actual Maturation | Predicted Maturation  (score) |
| --- | --- | --- | --- | --- |
| GSE70605#1 | Young | Young (10) | MII | MII (10) |
| GSE70605#2 | Young | Young (9.2) | MII | MII (10) |
| GSE70605#3 | Young | Young (9.5) | MII | MII (10) |

**Table S4 Sequencing of PCR primers**

| Gene | Primer | Sequence (5’-3’) |
| --- | --- | --- |
| *Gapdh* | F | GTGTTCCTACCCCCAATGTGT |
|  | R | ATTGTCATACCAGGAAATGAGCTT |
| *Ccnb1* | F | GCGTGTGCCTGTGACAGTTA |
|  | R | CCTAGCGTTTTTGCTTCCCTT |
| *Sycp3* | F | TGGAAACTCAGCAGCAAGAGA |
|  | R | GGCATGCCTCTTAGCTAATGTT |
| *Mos* | F | GATGTGGCTGGTTTTGAGAATC |
|  | R | AGGGAAGTTTGGGAGCATG |
| *Numb* | F | GCCGTAAAGAGATTGAAAGCTG |
|  | R | GTCTGGTCAACTATGAGGTCC |
| *Ezh2* | F | GCAACCCGAAAGGGCAAC |
|  | R | CCCACATACTTCAGGGCATCA |
| *Pou5f1* | F | CGTGGAGACTTTGCAGCCT |
|  | R | GCTTGGCAAACTGTTCTAGCTC |
